# Theta transcranial alternating current stimulation is not effective in improving working memory performance

**DOI:** 10.1101/2024.03.20.585954

**Authors:** Dauren Kasanov, Olga Dorogina, Faisal Mushtaq, Yuri G. Pavlov

## Abstract

There is an extensive body of research showing a significant relationship between frontal midline theta activity in the 4-8 Hz range and working memory (WM) performance. Transcranial alternating current stimulation (tACS) is recognized for inducing lasting changes in brain oscillatory activity. Across two experiments, we tested whether WM could be improved through tACS of dorsomedial prefrontal cortex and anterior cingulate cortex, by affecting executive control networks associated with frontal midline theta. In Experiment 1, following either a 20-minute verum or sham stimulation applied to Fpz-CPz at 1 mA and 6 Hz, 31 participants performed WM tasks, while EEG was recorded. The tasks required participants to either mentally manipulate memory items or retain them in memory as they were originally presented. No significant effects were observed in behavioral performance, and we found no change in theta activity during rest and task following stimulation. However, alpha activity during retention or manipulation of information in WM was less strongly enhanced during the delay period following verum stimulation as compared with sham. In Experiment 2 (N = 25), tACS was administered during the task in two separate sessions. Here, we changed the order of the stimulation blocks: a 25-minute task block was either accompanied first by sham stimulation and then by verum stimulation, or vice versa. Again, we found no improvements in WM through either tACS after-effects or online stimulation. Taken together, our results demonstrate that theta frequency tACS applied at the midline is not an effective method for enhancing WM.

## Introduction

Working memory (WM) is a crucial cognitive system, essential for temporarily storing and manipulating information, whether recently encountered or retrieved from long-term memory (Cowan, 2001; Miyake & Shah, 1999). Memorization, a key aspect of working memory, entails the encoding and storage of information, while manipulation involves the transformation of this stored information for diverse cognitive activities such as mental arithmetic, language comprehension, and spatial reasoning (Oberauer et al., 2000).

Neural oscillations in the theta band (4-8Hz), and specifically power enhancements in frontal midline theta, have been shown to be linked with WM performance across a range of tasks (Berger et al., 2019, 2014; Hsieh & Ranganath, 2014; Itthipuripat et al., 2013; Kawasaki et al., 2010; Pavlov & Kotchoubey, 2017, 2020, 2021). These theta oscillations are typically localized in the dorso-medial prefrontal cortex (dmPFC) and anterior cingulate cortex (ACC) (Holroyd & Umemoto, 2016). However, it is unclear whether the relationship between frontal midline theta, WM performance, and manipulations in WM is causal.

Transcranial alternating current stimulation (tACS) has the potential to entrain endogenous oscillatory brain activity and as such, help establish the causal relationship between brain rhythms and cognitive abilities (Thut et al., 2017). The application of theta tACS with the aim to improve WM has generated contradicting evidence. Some authors have concluded that theta tACS produced a significant improvement in WM scores (Biel et al., 2022; Hu et al., 2022; Vosskuhl et al., 2015; Wolinski et al., 2018; Zeng et al., 2022). In contrast, other have shown no significant impact (Abellaneda-Pérez et al., 2020; Feurra et al., 2016; Kleinert et al., 2017; Pahor & Jaušovec, 2018; Wolinski et al., 2018), or even disruptive effects (Chander et al., 2016) of the stimulation. This variability in outcomes may be attributed to methodological differences, including electrode placement, the participants’ state during stimulation, and the specific tasks chosen. We explore these possibilities in the present work.

The impact of tACS varies depending on whether stimulation occurs during task performance (referred to as ’online’) versus after the task (termed ’after-effects’). Research indicates that tACS after-effects can last from 30 minutes (Neuling et al., 2013) to 60 minutes (Wischnewski et al., 2019), and in some cases, up to 70 minutes (Kasten et al., 2016). However, these findings primarily relate to alpha tACS and may not apply to other frequency bands. A recent meta-analysis suggests that the after-effects of stimulation are often stronger than the online effects (Grover et al., 2023). Conversely, some studies (Pozdniakov et al., 2021) found that tACS influences cortical excitability only during online application, while others (Wischnewski et al., 2019) reported no significant difference between online and offline applications. In the domain of WM, a direct comparison showed no benefits of online stimulation but significant after-effects (Meiron & Lavidor, 2014; Vosskuhl et al., 2015). In contrast, others did not find any noticeable impact of tACS on behavior in either (Abellaneda-Pérez et al., 2020; Kleinert et al., 2017). Some studies employed a design with pre-, post-, and online assessments, which would allow the direct comparison of online and after-effects, however, they did not control for practice and fatigue effects in the later blocks of the task (Kleinert et al., 2017; Vosskuhl et al., 2015). Therefore, a more comprehensive comparison of online and after-effects of tACS is needed to better understand its potential for enhancing WM performance.

Theta stimulation targeting both the frontal-parietal region (Jaušovec et al., 2014a; Jones et al., 2019) and the parietal region alone (Jaušovec & Jaušovec, 2014) has proven more effective compared to stimulation that focuses solely on the frontal region (Jones et al., 2019; Jaušovec and Jaušovec, 2014). While electrodes in frontal-parietal montages are typically positioned in either the left or right areas, two studies placed the electrodes along the midline which produced the opposite effects - WM improvement (Vosskuhl et al., 2015) or deterioration (Chander et al., 2016). Midline electrode placements are theoretically more effective in targeting frontal midline theta sources - ACC and dmPFC. Therefore, in our study, we utilized an Fpz-CPz montage, which, according to simulations of electric current distribution, should effectively target both brain areas and circuits associated with the fronto-parietal executive control networks.

Another potential factor affecting efficacy of the stimulation is the ceiling effect, with chosen experimental tasks being too simple to attain additional benefits from the stimulation. This might mean that only participants with the lowest memory capacity could experience benefits from tACS. Consequently, some studies have reported improvements in WM only in a small group of participants with lower memory capacity scores (Reinhart & Nguyen, 2019; Zeng et al., 2022), and under specific conditions, such as in the most challenging tasks (Biel et al., 2022; Hu et al., 2022). To mitigate concerns regarding the ceiling effect, we incorporated tasks of varying difficulty levels in the present study.

Since frontal midline theta is more strongly involved in tasks engaging executive control networks, such as WM tasks requiring manipulations, the magnitude of the effect of the stimulation may be larger in such tasks. To date, the direct comparison of tACS effects between WM tasks requiring manipulations and simple retention of information in WM is inconclusive. For instance, Feurra et al. (2016) found no significant effects in both types of tasks: forward and backward digit span, while Vosskuhl et al. (2015) observed effects in the forward digit span task but not in the backward digit span task, and Jaušovec et al. (2014) reported beneficial effects in both tasks.

In the current study, we set out to test the utility of tACS as a tool for modulating WM processes. Here, our stimulation focussed on the ACC and dmPFC - shown to be sources of frontal midline theta activity appearing in WM tasks. We conducted two experiments using tasks that either required manipulation of WM content or merely the simple retention of memory items. We also adjusted the difficulty of these tasks to circumvent ceiling effects that could compromise the stimulation’s effectiveness. Building on the findings from the first experiment, we preregistered hypotheses for the second experiment to compare the online and after-effects of tACS.

### Experiment 1

We hypothesized that the after-effects of theta tACS would manifest in increased theta power during both rest and task periods, as well as in improved behavioral performance. We anticipated these effects to be more pronounced in tasks involving manipulations of WM content.

Our experimental design and analysis approach closely followed that of Berger et al. (2014), who employed verbal match-to-sample tasks aiming to differentiate between simple retention of information in WM and mental manipulation of WM content with and without involvement of access to long-term memory. We aimed to replicate the effects of task type on theta, alpha, and beta activity during the delay period. In addition to varying the type of task, we also varied the WM load between 4 and 6 letters, thereby extending the original study. Consequently, a secondary goal of Experiment 1 was to investigate oscillatory brain activity (1) during WM manipulations that either require access to long-term memory or only use sensory content as compared with simple retention, and (2) under conditions of varying WM load.

#### Experiment 1: Methods

##### Participants

Data were collected from 31 participants (15 female; M_age_ = 24.3, SD_age_ = 7.05). All but two were right handed (Annett, 1970). Before inclusion the participants were screened to not have any metal or electronic implants, cardiostimulators, pregnancy, and epilepsy. They had been free of using any medication, had not had any psychiatric/neurological conditions (e.g., depression, bipolar disorder, epilepsy, migraine, severe head trauma, brain surgery) currently or in the past. They had normal or corrected to normal vision. All participants gave written informed consent. The experimental protocol for both experiments was approved by the ethics committee of Ural Federal University.

The sample size was based on previous similar studies. For comparison, tACS studies on working memory that employed a within-subject design, as reported in a recent meta-analysis (Grover et al., 2023), had a median sample size of 17.

Six EEG recordings were excluded due to data loss, two more were excluded due to excessive amounts of artifacts (less than 12 clean trials in any of the conditions). Thus, 23 participants were included in the final EEG analysis.

##### Task

We used a set of delayed match-to-sample WM tasks (Figure 1). Participants were presented with strings of four or six capital letters from the Russian alphabet (only consonants). Each string was displayed for two seconds, followed by a delay period of 6.5 seconds. After this delay, a probe letter string was shown for one second. The participants indicated whether the probe letters matched the original set (or the set after mental manipulations) or differed from it, during or after the probe’s appearance, by pressing a button on a gamepad. The participants were instructed to provide the most accurate answers possible, but were warned that time was limited. The time to respond after the probe’s offset was limited to 5 seconds. The likelihood of the probe being a mismatch was 50% with only one letter altered in each case. The inter-trial interval randomly varied between 5 and 5.5 seconds. Psychtoolbox (Kleiner et al., 2007) was used for the stimulus presentation.

There were three types of the task, each cued by a specific instruction before the letters were shown: “Forward,” “Backward”, and “Alphabetical”. Two types of the task required manipulation with the memory content during the delay period. In the “Backward” manipulation task, participants were asked to mentally reverse the order of the letters in memory. In the “Alphabetical” manipulation task, they were required to mentally rearrange the letters in alphabetical order. The “Forward” task, on the other hand, simply involved retaining the letter set as presented, without any mental reordering. The task instruction, letters for encoding, and the probe had the following font color: Forward - green, Alphabetical - red, Backward - blue (see Figure 1).

There were 120 trials in total with 20 trials per condition. The trials were presented in blocks of 10 corresponding to one of the 6 possible conditions. The duration of the task was ∼40 min.

For each participant, proportions of hits and false alarms were used to compute a sensitivity index d’ as the difference in standardized normal deviates of hits minus false alarms (Böckmann-Barthel, 2023). Perfect rates were corrected according to 1/2N rule (Hautus, 1995). d’ served as our main outcome measure of the behavioral performance, however, we also report reaction time (RT) for completeness. For RT analyses, at the level of an individual participant, we first calculated the median value and then, in the group analyses, we used the mean. All responses, correct and incorrect, were included in the analysis. The average accuracy in percentage of correct responses was also calculated for each type of task.

**Figure 1.**
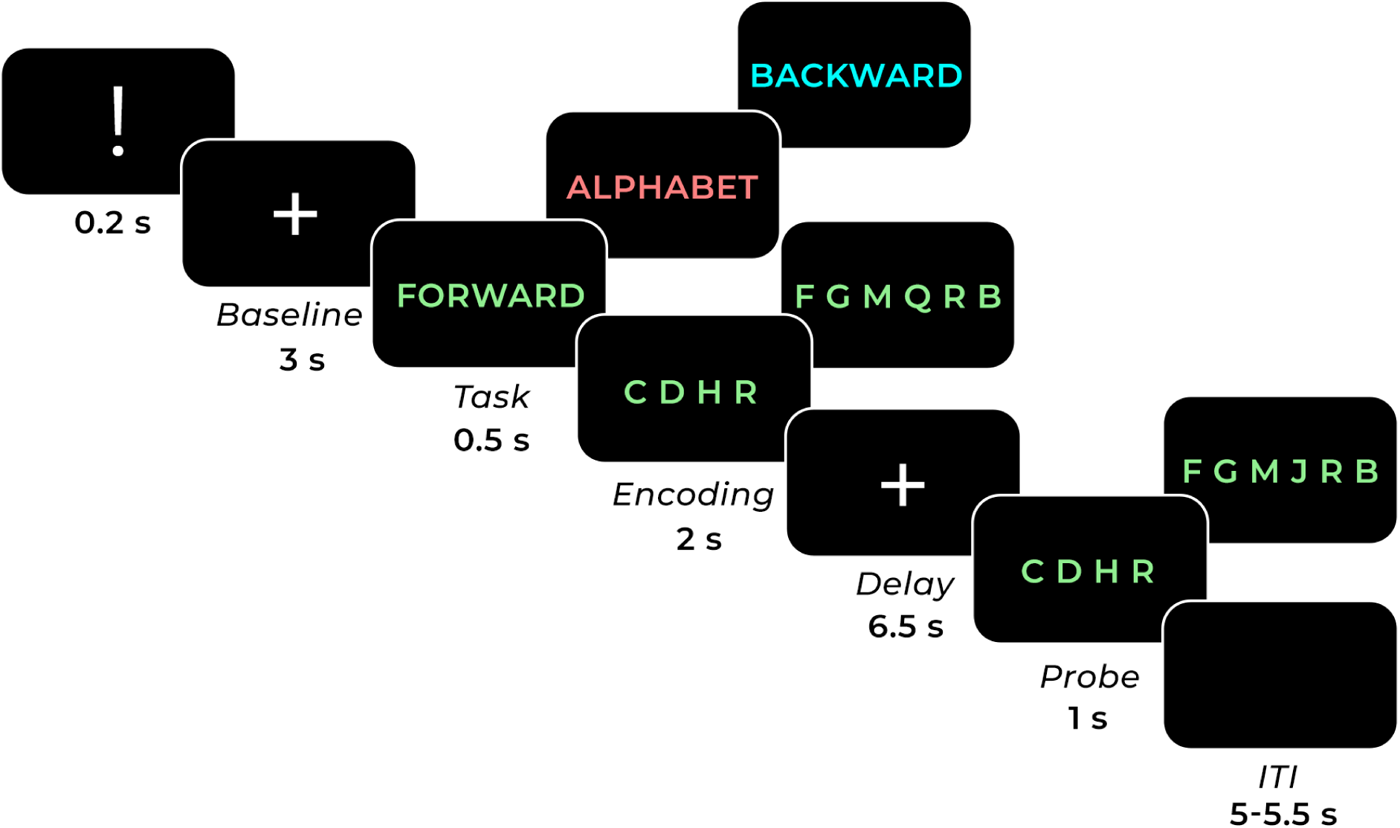
Visual representation of the experimental paradigm (not to scale). In each trial, participants memorized a string of letters (with a length of 4 or 6) that were presented simultaneously. The type of task varied between trials: participants were required to either memorize the letters as they were presented (in forward order), mentally reorder them in reverse sequence during the delay period (backward order), or mentally rearrange the letters in alphabetical order. After the delay, a probe appeared, and participants pressed a button to indicate whether the probe string matched the original string (in the case of forward order) or matched the string after mental manipulation (in the cases of backward and alphabetical order).

##### Brain stimulation

The participants completed two sessions (average time between 1 and 2 sessions was 7.06 days; range 4-15 days). The order of the sham and verum sessions was randomly assigned for each participant.

At the beginning of each session, we set up the stimulation electrodes and checked the subelectrode impedance to ensure it was maintained below 10 kΩ. The amplitude of the stimulating current was based on the thresholds individually determined for skin sensations induced by the stimulation. During the calibration, tACS was applied with the parameters described below for 10 seconds. Participants were then asked about their tolerance to the stimulation, specifically inquiring about skin sensations, phosphenes, or any discomfort they might experience. If they reported discomfort that could prevent their participation for the entire session, we decreased the amplitude in steps of 0.05 mA. Conversely, if they did not feel the stimulation tickling sensation, we increased the amplitude. The calibration procedure resulted in a peak-to-peak amplitude of 1.25 mA in two cases, 1.5 mA in three cases, and 1 mA for the remainder.

tACS was applied over Fpz and CPz at 6 Hz with standard 7 x 5 electrodes covered by sponges soaked with saline. Verum stimulation had the following parameters: 8 seconds acceleration, 20 minutes tACS, 5 seconds deceleration, while sham was different in duration of the tACS and ended after 30 seconds of stimulation. For transcranial brain stimulation we used neuroConn DC-STIMULATOR PLUS. The location of stimulation electrodes was based on the results of a simulation of electric field distribution performed in SimNIBS 3.2.4 to target the dmPFC and ACC regions (see Figure 2).

**Figure 2.**
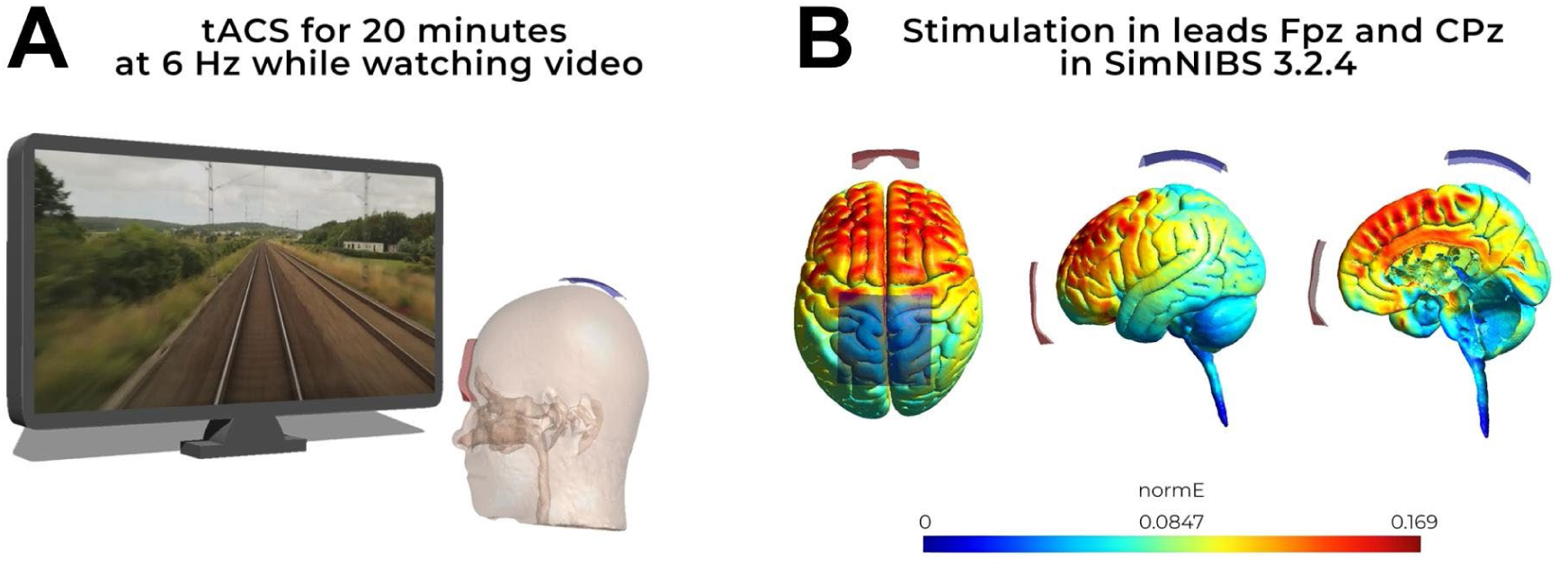
(a) The stimulation session lasted for 20 minutes. During this period, participants viewed a video on the screen depicting train movement, recorded from the driver’s cabin. (b) Location of the stimulation electrodes and the results of the simulation of the currents. normE - electric field intensity distribution, V/m^2^.

During the stimulation, participants watched a video recording of a train from the cabin’s perspective. The lights were on. Following stimulation, the participants self-assessed severity of the symptoms (1 - none, 2 - mild, 3 - moderate, 4 - strong): headache, neck pain, sore scalp, scalp burns, skin tingling, skin burning, drowsiness, difficulty concentrating, mood swings, other; and the association of symptoms with stimulation (1 - no, 2 - unlikely, 3 - possible, 4 - definitely).

Directly after the stimulation, the EEG capping procedure was initiated. It led to a delayed EEG recording onset with the following delays (mean ± SD, min): resting state, verum: 18.7 ± 11.2, sham: 22 ± 14.2; WM task, verum: 33.7 ± 17.7, sham: 36.2 ± 14.3.

At the end of the final session, we assessed the effectiveness of blinding by asking participants to identify which sessions they believed involved real stimulation, with possible answers being sham or verum for each session. This created four possible response combinations: verum/verum, sham/sham, sham/verum, and verum/sham. We then performed a paired samples t-test to compare the responses between the sham and verum sessions. The results indicated that participants were unable to accurately identify the type of stimulation they received (t(30) = 0.94, p = 0.354).

##### Electroencephalography

EEG data were recorded with a 129-channel Geodesic Sensor Net. EEG was online referenced to Cz and digitized continuously at 1000 Hz. Impedances were maintained below 50 kΩ as recommended to ensure an optimal signal-to-noise ratio for this amplifier system (NetAmps 300). During EEG recording, the lights were off; if necessary, forced ventilation was turned on. Before the WM task, resting state EEG with eyes closed was recorded for 3 minutes and with eyes open for 1.5 minutes.

EEGLAB (Delorme & Makeig, 2004) was used for data preprocessing. Each recording was filtered by applying 1 Hz high-pass and 45 Hz low-pass filters (pop_eegfiltnew function in EEGLAB). Then, the data were downsampled to 250 Hz and bad channels (containing more than 20% of artifacts) were visually identified and restored by means of spherical interpolation. An Independent Component Analysis (ICA) was performed using the AMICA algorithm (Palmer et al., 2012). We first used ICLabel (Pion-Tonachini et al., 2019) to remove components which were classified with probability to be brain activity less than 5% or labeled otherwise with more than 80% probability. Remaining components clearly related to eye movements and high amplitude muscle activity were removed manually. After that, the data were re-referenced to average reference.

For WM task data analysis, the data were epoched into [-7000 8000 ms] intervals with 0 representing onset of the retention interval. In epochs containing up to 5 bad channels, the channels were spherically interpolated. Remaining epochs still containing artifacts were visually identified and discarded. All epochs were then converted into current source density (CSD) using CSD toolbox (Kayser, 2009). We used spherical spline surface Laplacian (Perrin et al., 1989) with the following parameters: 50 iterations; m = 4; smoothing constant λ = 10^−5^ (Tenke & Kayser, 2005).

Time-frequency analysis was performed on the preprocessed single trial data between 1 and 45 Hz with 1 Hz steps using Morlet wavelets with the number of cycles varying from 3 to 12 in 45 logarithmically spaced steps for each participant and condition, separately. The analysis time window was shifted in steps of 20 ms. Spectral power was baseline-normalized by computing the percent change of the power with respect to the [-3700 to -2700 ms] time interval, which corresponded to part of the baseline fixation before the presentation of the task instruction. The time-frequency analysis was performed by means of the Fieldtrip toolbox (Oostenveld et al., 2011).

For resting state analysis, preprocessing steps of the time interval corresponding to the eye-closed state were the same as for the WM task data. The data were divided into epochs of 2 seconds, converted into CSD, and the Fourier transform was carried out with a Hanning window with a frequency resolution of 0.5 Hz. For the direct comparison and visualization, the task data corresponding to 0.5-6.5 s time interval where 0 is the onset of the delay period were additionally analyzed in the same way as the resting state data.

Three frequency bands and regions of interest (ROI) were identified: frontal theta (4-8 Hz, channels E15, E18, E10, E11, E16), posterior alpha (8-13 Hz, channels E60, E62, E85, E59, E67, E77, E91, E58, E66, E72, E84, E96, E65, E90, E70, E75, E83), and beta (13-20 Hz, the same channels as for alpha). The theta channels were selected because they are located around Fz channel, which has been used for quantifying frontal midline theta in most previous studies (Pavlov & Kotchoubey, 2020). Alpha ROI covers a large part of the occipito-parietal cortex where alpha enhancement during the delay period was visually strongly present on the flatten average (Bowman et al., 2020). Following the analysis protocol of Berger et al. (2014), we selected posterior channels to quantify beta activity, same channels as for alpha. The same frequency bands and ROIs used in the analysis of the resting state data.

##### Statistics

To test the effects of tACS on behavioral performance, we used a repeated-measures ANOVA with the factors Task (Backward manipulation, Alphabetical manipulation, Retention), Load (4 or 6 letters), and Stimulation (Sham vs Verum). Greenhouse-Geisser adjusted degrees of freedom are reported where applicable. The analysis was repeated for d’ and RT.

The same analysis was used for testing the effects of Stimulation, Task, and Load on relative spectral power in theta, alpha, and beta frequency bands.

We tested the lack of difference between stimulation conditions by means of Bayesian ANOVA with default priors (Cauchy(scale=sqrt(2)/2)) using BayesFactor library (v. 0.9.12.4.2) for R (Morey & Rouder, 2018). We compared the models with inclusion of the Stimulation effect against all other models using the bayestestR::bayesfactor_inclusion function. Inclusion (or exclusion) Bayes factors are reported unless specified otherwise.

Statistical analyses in both experiments were conducted using R (v. 4.2.2).

#### Experiment 1: Results

##### Behavior

We examined how theta tACS affects d’ and RT in WM tasks. These tasks either mainly depend on sensory components, like memorizing sets of 4 or 6 letters in a forward order (the retention task), or heavily involve executive components, such as alphabetical and backward manipulation tasks. We anticipated that the effect would be more pronounced in the manipulation tasks than in the retention task.

As expected, performance was worse in the 6-letter as compared with 4-letter condition (main effect of Load; see Table 1 and Figure 3). The type of task significantly affected behavioral performance with lowest d’ attained in the alphabetical manipulation condition, highest in the retention condition, and the backward manipulation condition in the middle (main effect of Task; backward < alphabetical: t(30) = 7.85, p < 0.001, dz = 1.43; backward < retention: t(30) = 4.77, p < 0.001, dz = 0.87; alphabetical < retention: t(30) = 10.6, p < 0.001, dz = 1.93). No significant interactions of Load and Task were found. The stimulation did not produce any significant effects or interactions. The lack of the stimulation effect was confirmed by the Bayesian ANOVA showing support for the null hypothesis (the effect of Stimulation on d’: exclusion BF = 8.93).

**Table 1.** The result of ANOVA for d’.

| Effect | df | F | $\eta_p^2$ | p |
| --- | --- | --- | --- | --- |
| Load | 1, 30 | 199.82 | 0.869 | <.001 |
| Stimulation | 1, 30 | 1.55 | 0.049 | 0.222 |
| Task | 1.47, 44.07 | 78.07 | 0.722 | <.001 |
| Load x Stimulation | 1, 30 | 0.11 | 0.004 | 0.744 |
| Load x Task | 1.75, 52.63 | 2.16 | 0.067 | 0.131 |
| Stimulation x Task | 1.71, 51.24 | 0.29 | 0.010 | 0.712 |
| Load x Stimulation x Task | 2, 59.97 | 2.59 | 0.080 | 0.083 |

**Figure 3.**
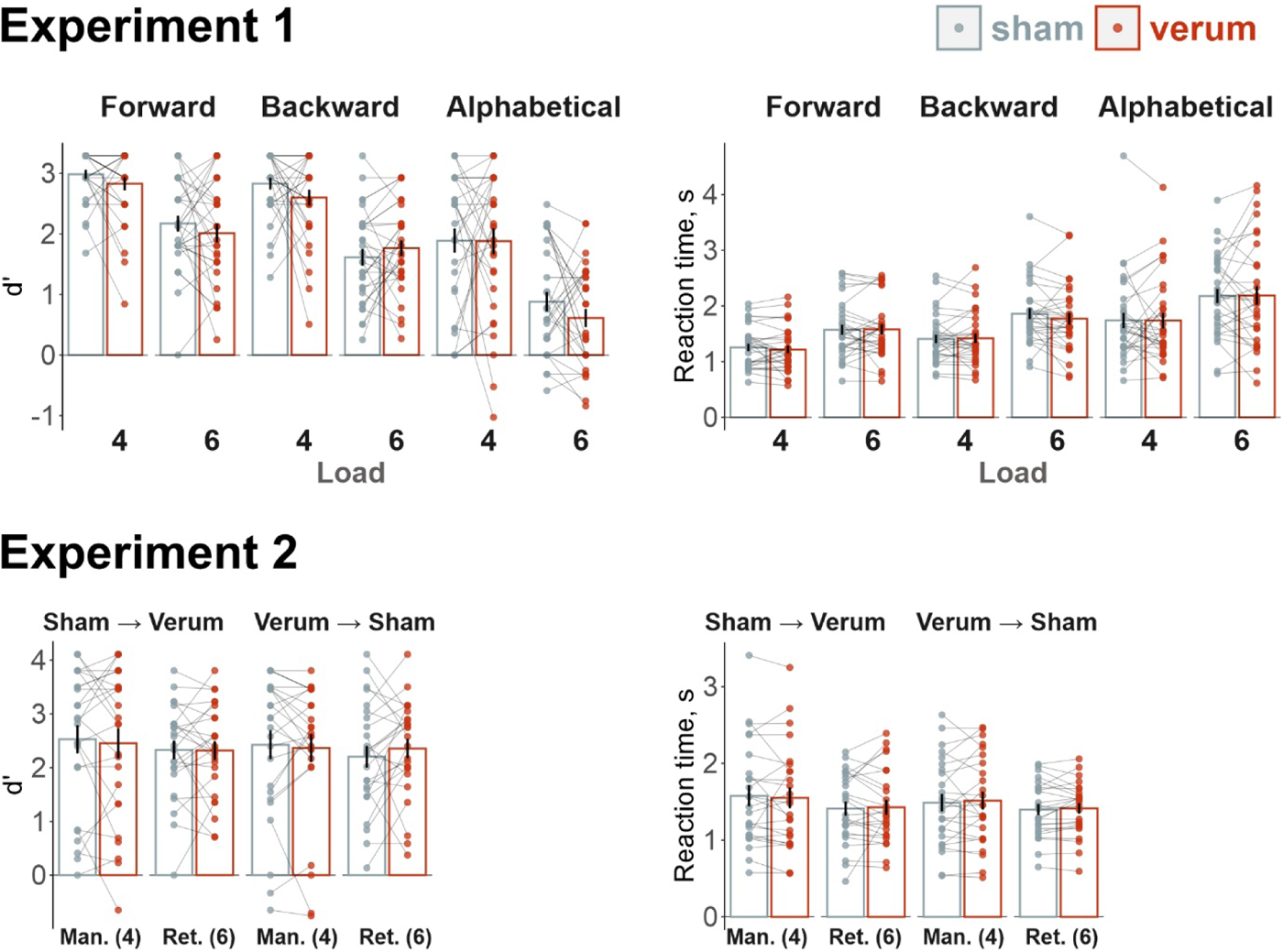
Results of the behavioral data analyses. In Experiment 2, Verum -> Sham and Sham -> Verum indicate order of the stimulation blocks. Man(4) is the alphabetical manipulation of 4 letters condition, Ret(6) is the retention of 6 letters condition. Error bars are the standard error of the mean.

The ANOVA results for RT mirrored those obtained for d’ (see Figure 3 and Table 2). Participants responded more quickly in the 4-letter condition compared to the 6-letter condition (main effect of Load). The RT pattern with retention < backwards < alphabetical was also confirmed (main effect of Task; backward < alphabetical: t(30) = 3.56, p = 0.003, dz = 0.65; backward < retention: t(30) = 6.64, p < 0.001, dz = 1.21; alphabetical < retention: t(30) = 6.01, p < 0.001, dz = 1.1). There was no significant interaction between Load and Task. Additionally, we attempted to replicate the effects of theta tACS on RT in the task of retaining 6 letters among high-performing individuals (identified using a median split based on overall accuracy), as previously demonstrated by Hu et al. (2022). This analysis found no significant effects (low-performance group: t(15) = 0.61, p = 0.559, dz = 0.158; high-performance group: t(14) = 0.96, p = 0.352, dz = 0.257). Finally, stimulation did not produce any significant effects or interactions.

**Table 2.** ANOVA for RT.

| Effect | df | F | $\eta_p^2$ | p |
| --- | --- | --- | --- | --- |
| Load | 1, 30 | 65.64 | 0.686 | <.001 |
| Stimulation | 1, 30 | 0.13 | 0.004 | 0.722 |
| Task | 1.16, 34.91 | 24.84 | 0.453 | <.001 |
| Load x Stimulation | 1, 30 | 0.06 | 0.002 | 0.814 |
| Load x Task | 1.37, 41.02 | 0.82 | 0.027 | 0.406 |
| Stimulation x Task | 1.24, 37.22 | 0.21 | 0.007 | 0.702 |
| Load x Stimulation x Task | 1.39, 41.74 | 0.91 | 0.029 | 0.379 |

##### EEG

To determine whether tACS produces any significant impact on brain activity, we explored EEG during the resting state and task performance across three frequency bands: theta, alpha, and beta. Additionally, a deeper exploration of the task (retention, alphabetical, or backward manipulation) and load (4 or 6 letters) effects allowed us to test the replicability of the results obtained by Berger et al. (2014).

In the resting state, theta activity was not significantly different between the stimulation conditions (F(1, 24) = 1.20, p = .285, *η_p_^2^* = .05; Figure 4a). Similarly, alpha power was not significantly affected by the stimulation (F(1, 24) = 0.55, p = .466, *η_p_^2^* = .02).

In the WM task, theta activity during the delay period (0.5-6.5 s) was more pronounced in the 6-letter condition compared to the 4-letter conditions (main effect of Load; see Table 3 and Figure 5). Theta power was highest in the alphabetical order condition, followed by the backward order, and finally, the forward order condition, with all pairwise comparisons showing statistical significance (p < 0.05). However, no effects of stimulation in the theta band were observed during the task (see Figure 4b).

**Table 3.** ANOVA for theta over the frontal ROI.

| Effect | df | F | $\eta_p^2$ | p |
| --- | --- | --- | --- | --- |
| Task | 1.48, 32.6 | 8.04 | 0.268 | 0.003 |
| Load | 1, 22 | 4.84 | 0.180 | 0.039 |
| Stimulation | 1, 22 | 0.60 | 0.027 | 0.446 |
| Task x Load | 1.4, 30.73 | 3.42 | 0.134 | 0.061 |
| Task x Stimulation | 1.74, 38.2 | 0.54 | 0.024 | 0.564 |
| Load x Stimulation | 1, 22 | 0.40 | 0.018 | 0.531 |
| Task x Load x Stimulation | 1.69, 37.07 | 0.19 | 0.008 | 0.793 |

**Figure 4.**
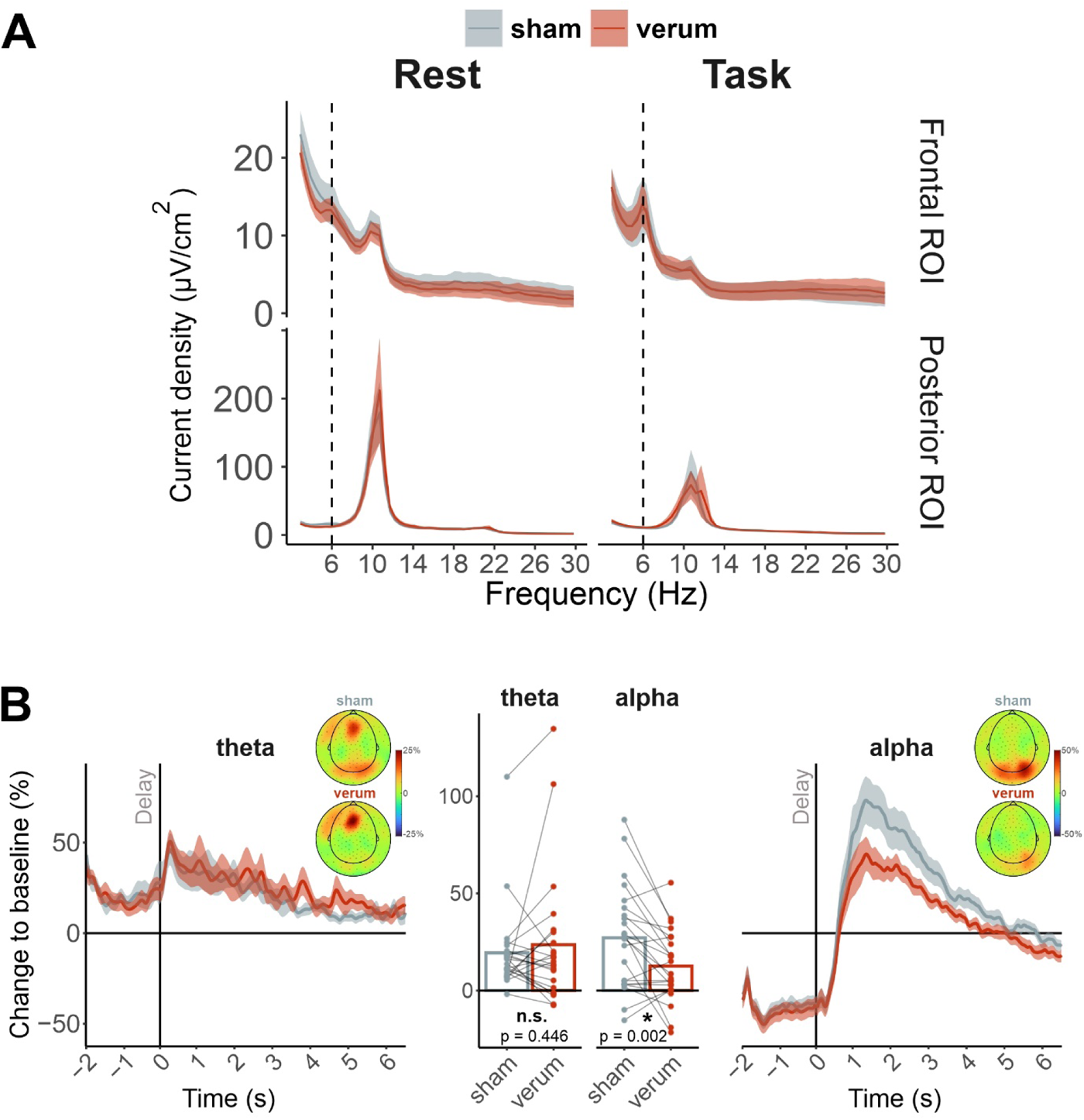
Stimulation effects on EEG. (a) Distribution of absolute values of current source density in the resting state and WM task (averaged over 0.5-6.5 s after the delay onset) over the frontal and posterior ROIs. (b) Comparison of baseline normalized current source density between Sham and Verum stimulation conditions in theta and alpha frequency bands. The average theta activity during the delay period (0.5-6.5 s time window) over the frontal ROI did not differ between stimulation conditions, whereas alpha activity over the posterior ROI was significantly stronger after sham stimulation.

**Figure 5.**
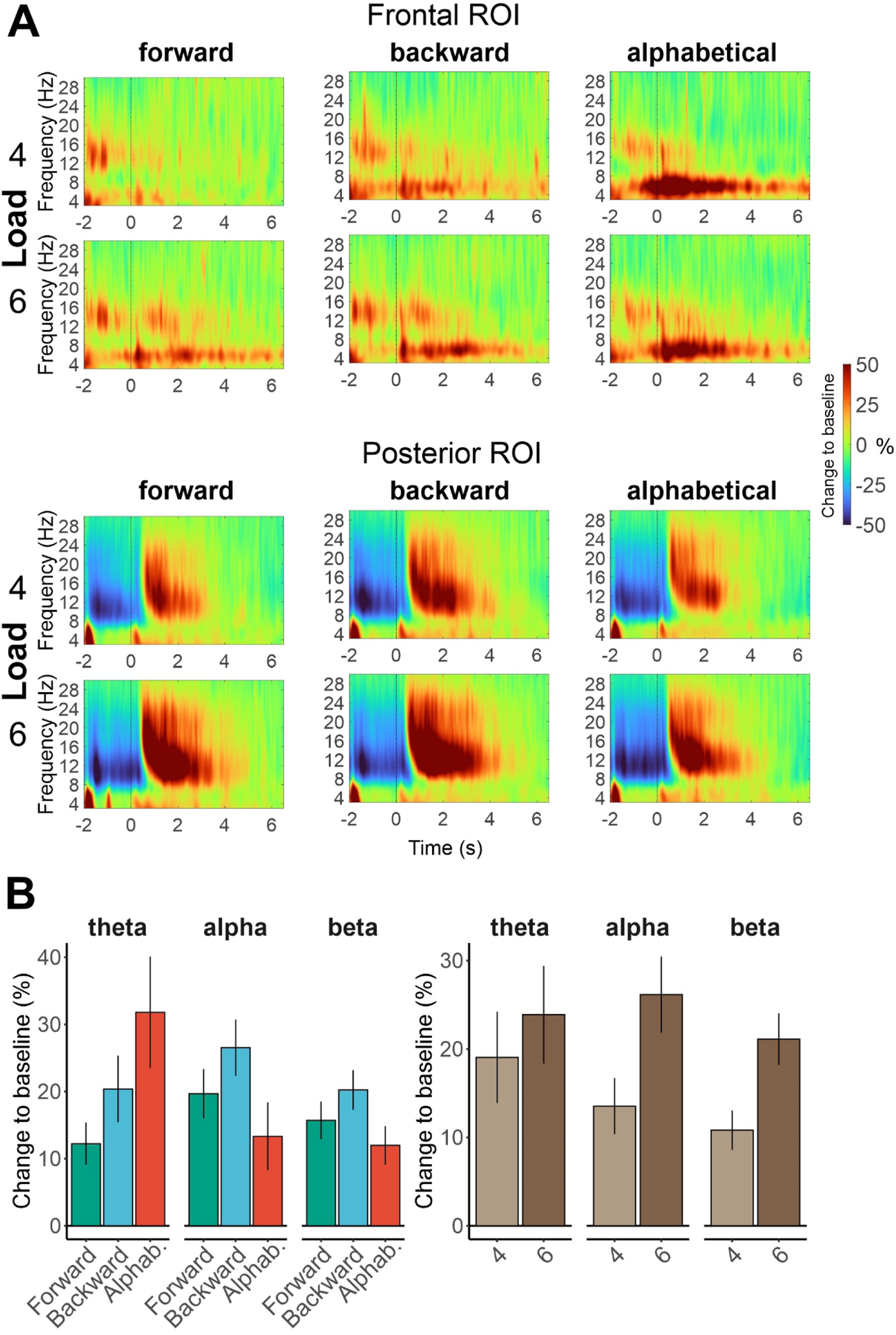
Results of the time-frequency analysis of general effects of Task and Load (a) Time-frequency maps showing the effects of Task and Load on spectral power (baseline normalized current source density). (b) Comparison of the mean values between the conditions averaged over the delay period (0.5-6.5 s time window). Theta activity over the frontal ROI and alpha activity over the posterior ROI increased with WM load. Theta showed the strongest increase in the most challenging condition - the alphabetical manipulation. For alpha, alphabetical manipulation resulted in the smallest increase. Error bars represent standard error of the mean.

The increase in alpha activity during the delay period was larger in the 6-letter condition compared to the 4-letter condition, indicating a main effect of Load (see Figure 5 and Table 4). Furthermore, alpha activity was highest in the backwards manipulation condition, followed by the retention and then the alphabetical manipulation condition (main effect of Task). In the post-hoc t-tests, only the backwards and alphabetical manipulation tasks showed a significant difference (p < 0.05). Baseline-normalized alpha activity over posterior channels appeared stronger after sham stimulation compared to verum. This effect was not influenced by the factors of Load or Task.

**Table 4.** ANOVA for alpha activity over the posterior ROI.

| Effect | df | F | $\eta_p^2$ | p |
| --- | --- | --- | --- | --- |
| Task | 1.82, 40.1 | 4.16 | 0.159 | 0.026 |
| Load | 1, 22 | 19.05 | 0.464 | <.001 |
| Stimulation | 1, 22 | 11.97 | 0.352 | 0.002 |
| Task x Load | 1.91, 41.99 | 0.57 | 0.025 | 0.562 |
| Task x Stimulation | 1.94, 42.75 | 0.58 | 0.026 | 0.56 |
| Load x Stimulation | 1, 22 | 0.01 | <.001 | 0.926 |
| Task x Load x Stimulation | 1.87, 41.21 | 1.34 | 0.057 | 0.272 |

The effects in the beta frequency band were identical to the ones described above for alpha (Table 5).

**Table 5.** ANOVA for beta over the posterior ROI.

| Effect | df | F | $\eta_p^2$ | p |
| --- | --- | --- | --- | --- |
| Task | 1.87, 41.2 | 4.19 | 0.160 | 0.024 |
| Load | 1, 22 | 27.14 | 0.552 | <.001 |
| Stimulation | 1, 22 | 6.98 | 0.241 | 0.015 |
| Task x Load | 1.9, 41.9 | 3.03 | 0.121 | 0.061 |
| Task x Stimulation | 1.83, 40.29 | 0.83 | 0.036 | 0.434 |
| Load x Stimulation | 1, 22 | 0.64 | 0.028 | 0.433 |
| Task x Load x Stimulation | 1.97, 43.39 | 0.72 | 0.032 | 0.49 |

### Experiment 2

Experiment 1 found no significant effects of tACS on behavioral performance. We hypothesised that this may be due to at least three factors: (1) the task was administered with a ∼ 30 min delay after the tACS application, potentially diminishing any positive effects; (2) the stimulation occurred before, rather than during, the task, which may lessen the impact of the stimulation as online effects could be more pronounced; (3) the experiment consisted of only one stimulation session, which might not be sufficient. Additional stimulation in subsequent sessions could accumulate effects, thereby intensifying them. To address these potential explanations, we conducted a follow up study on behavioural performance and pre-registered (see https://osf.io/9gnhv) the following predictions:

1. tACS during WM task (online stimulation) positively affects behavioral performance;
2. There is an after-effect of online tACS: Performance in the task block following online tACS will be superior compared to that following sham stimulation;
3. The after-effect of tACS is less pronounced than the effect of online stimulation;
4. The effect of tACS is stronger in the manipulation task than in the simple retention task.

These predictions are visually depicted in Figure 6 (left panel). To preview the results (Figure 6, right panel), none of our predictions were confirmed.

**Figure 6.**
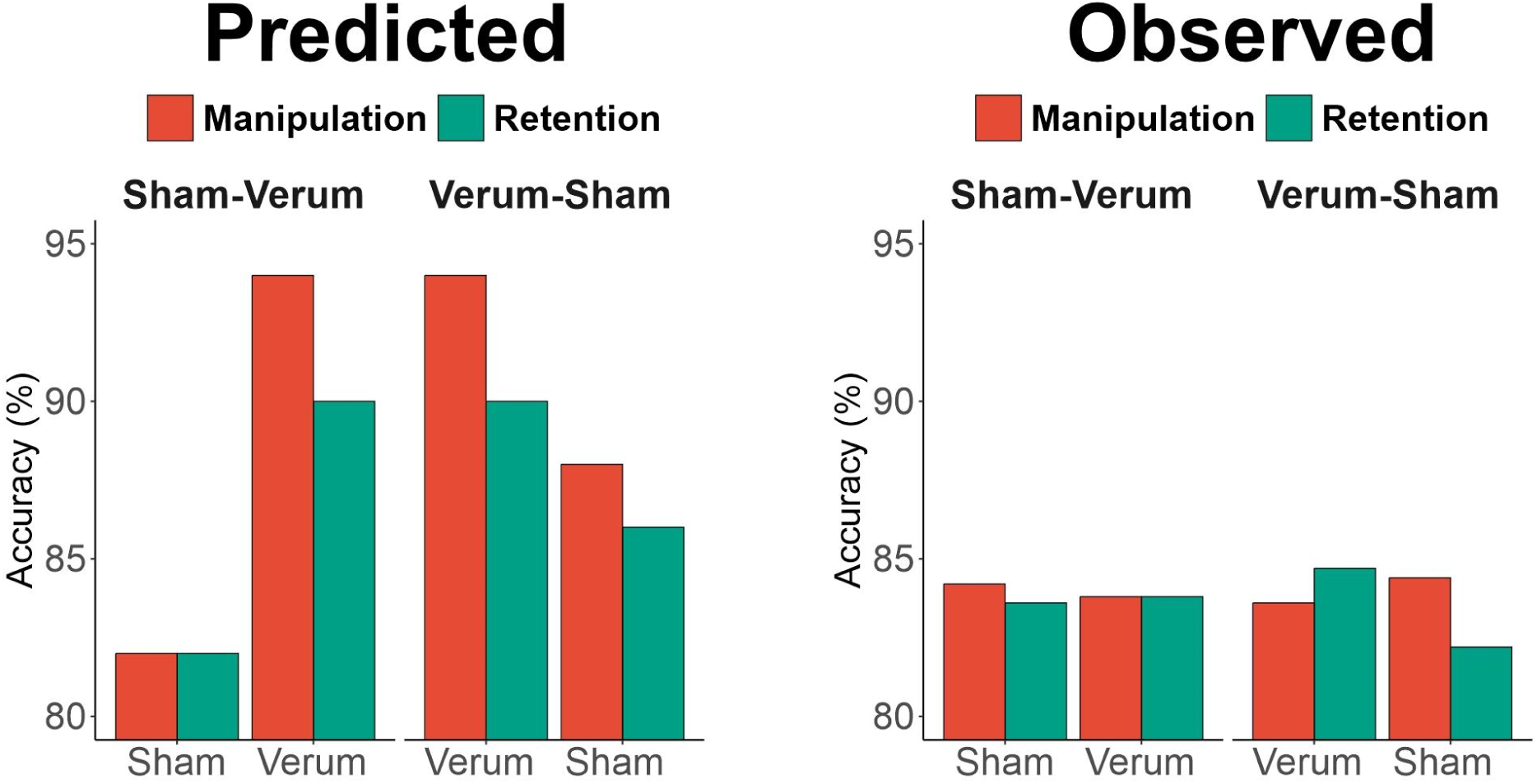
Difference between Predicted and Observed Results for Experiment 2. Left panel: Hypothetical distribution of accuracy in Experiment 2. Verum-Sham and Sham-Verum indicate order of the stimulation blocks in the corresponding sessions. Right panel: Observed distribution of accuracy in Experiment 2. Note that our main outcome variable is d’, but the distribution is shown for accuracy. The results for d’ and RT are shown in Figure 3.

#### Experiment 2: Methods

##### Participants

After screening using the same criteria as in Experiment 1, a group of 27 participants were recruited from the local student population of Ural Federal University and by online advertisements in the social media. Twenty-five of them participated in all three sessions and were included in the analysis (18 females; M_age_ = 20.9, SD_age_ = 4.21). They had normal or corrected to normal vision and reported to have no prior neurological conditions, and normal intelligence according to Raven’s Progressive Matrices. All but four were right-handed.

Sample size was determined using power analysis. Our goal was to obtain 0.8 power to detect a medium effect size of Cohen’s d = 0.5. Power analysis in G*Power revealed that with 25 participants we would be able to detect an effect with Cohen’s d = 0.48.

##### Task and procedure

The WM task was essentially the same as in Experiment 1. Briefly, participants were instructed to memorize strings of either 6 letters without any manipulation as they were presented (retention task) or 4 letters after mental recombination of letters in the alphabetic order (manipulation task). We made some changes to the task based on the results from Experiment 1. First, Experiment 1 showed equivalent performance in the retention of six letters and alphabetical manipulation of four letters and as such we retained only these two conditions. Second, we implemented several changes to reduce the experiment’s duration: (1) the delay period was shortened to six seconds; (2) the response time window after probe onset was limited to six seconds, after which the next trial began automatically; (3) the inter-trial interval (ITI) depended on reaction time - the reaction time was subtracted from a maximum of five seconds for ITI; (4) the fixation period before presenting the task instruction was omitted.

Five similar 25-min blocks filled with the same working memory task were distributed over 3 sessions, one per day. Each block consisted of 50 trials of both types (i.e., manipulation and retention tasks). The trials were presented in groups of 25 corresponding to the same type of the task (i.e., retention or manipulation). The type of task switched sequentially after every 25 trials. The first trial’s type was randomly assigned.

The same group of participants was assessed in all three sessions. The average time between the first and second sessions was 8.8 days (range: 1-25 days), and the time between the second and third sessions was 8.76 days (range: 6-23 days). The first session was always a training session, during which no stimulation electrodes were applied. Participants completed demographic forms, practiced a block of the task, and took the Raven Progressive Matrices test. In the second session (SHAM-VERUM), participants completed a brief practice (8 trials) and two full blocks of trials with a break in between. During the first block, sham stimulation was applied, and during the second block, verum stimulation was applied. Consequently, there were no after-effects of stimulation during the second block. In the third session (VERUM-SHAM), the procedure was identical, except that the first block was accompanied by tACS, and the second block by sham stimulation. In this case, we expected the after-effects of tACS to occur during the sham block. The order of the SHAM-VERUM and VERUM-SHAM sessions was randomized between participants. Only data from the second and third days were included in the analyses.

**Figure 7.**
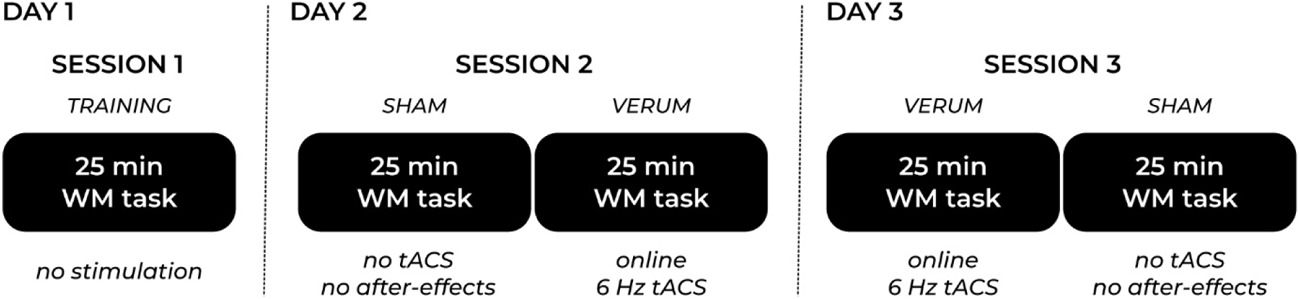
Design of Experiment 2. On the first day, participants completed a 25-minute block of WM task containing 25 manipulation and 25 retention trials. On the second and third days, the order of 25- min long blocks of the task (identical to the training session), but the task was administered either during sham or verum stimulation. The order of Sham-Verum and Verum-Sham sessions was randomized between participants.

##### Brain stimulation

The stimulation parameters were identical to Experiment 1, except stimulation lasted 25 minutes in the verum blocks. Calibration of individual stimulation amplitude was the same as in Experiment 1. The stimulation amplitude was adjusted in 6 participants: in 3 participants it was decreased to 0.95, 0.750 and 0.65 mA, in the other 3 participants it was increased to 1.2, 1.25, and 1.25 mA. For all other participants it remained exactly at the level of 1 mA peak-to-peak.

##### Statistics

For both d’ and RT, we ran an ANOVA with 3 within-subject factors: Task (Manipulation vs Retention), Order (Sham-Verum vs Verum-Sham), and Stimulation (Sham vs Verum).

#### Experiment 2: Results

The ANOVAs conducted to evaluate the impact of tACS on behavioral performance indicated no significant effects (for d’ see Table 6; for RT see Table 7). The Bayesian ANOVA showed evidence for the null hypothesis (the effect of Stimulation on d’: exclusion BF = 17.2; RT: exclusion BF = 16.4). Furthermore, we investigated whether a cumulative effect of stimulation was present, which would be evidenced by more pronounced online effects on the third day compared to the second day of the study. However, no significant effects were observed in this regard as well.

**Table 6.**
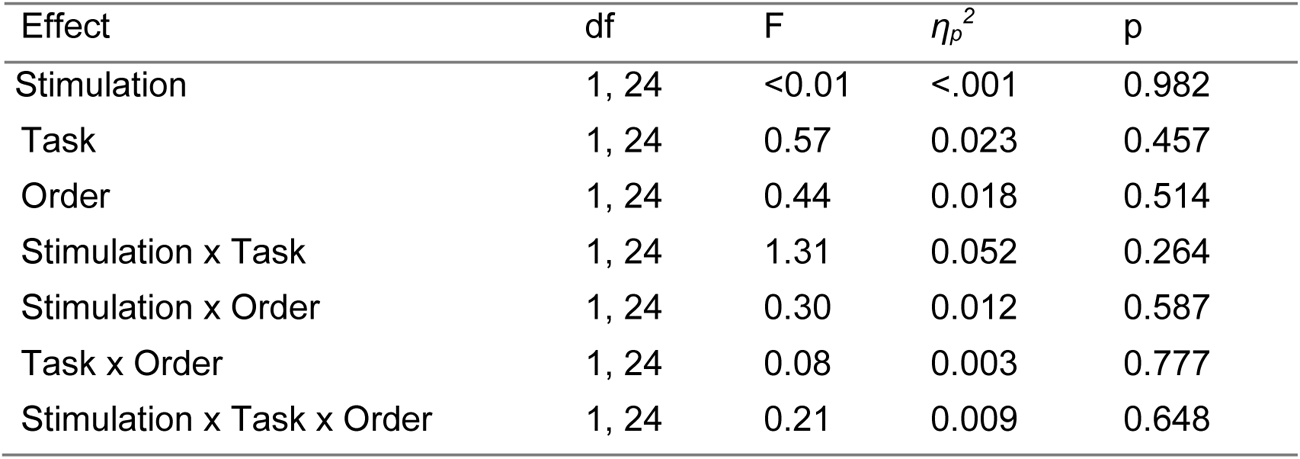
ANOVA for d’.

**Table 7.** ANOVA for RT.

| Effect | df | F | $\eta_p^2$ | p |
| --- | --- | --- | --- | --- |
| Stimulation | 1, 24 | 0.11 | 0.005 | 0.743 |
| Task | 1, 24 | 2.44 | 0.092 | 0.131 |
| Order | 1, 24 | 0.51 | 0.021 | 0.482 |
| Stimulation x Task | 1, 24 | 0.20 | 0.008 | 0.656 |
| Stimulation x Order | 1, 24 | 0.13 | 0.005 | 0.722 |
| Task x Order | 1, 24 | 0.98 | 0.039 | 0.333 |
| Stimulation x Task x Order | 1, 24 | 0.46 | 0.019 | 0.502 |

## Discussion

We tested the effect of stimulation of dmPFC and ACC at theta frequency on working memory (WM) performance in tasks that either strongly engage executive components (manipulation tasks) or rely primarily on sensory components (retention task) of WM. We expected the effect to be stronger in the manipulation task than in the retention task. In contrast to our expectations, we found that neither online stimulation during performance of the task nor the stimulation applied before the task execution produces any noticeable effects on measures of accuracy or reaction time in both types of the task.

### Why did theta tACS not affect WM performance?

Several factors might have affected the null findings in our study, and we reflect on possible explanations next.

We hypothesized that targeting frontal midline theta sources – ACC and dmPFC – would produce the strongest effects on WM. Previous research had shown success with more posterior and lateral electrode placements, such as the F4-P4 montage or targeting the bilateral parietal cortex (P3-P4 montage) (Jaušovec & Jaušovec, 2014; Jaušovec et al., 2014b; Jones et al., 2019). Our stimulation montage, however, might have been less effective in synchronizing frontal and posterior brain areas within the executive control network. Considering the role of brain oscillations in facilitating communication between distant brain areas – specifically sensory and executive control hubs – it is possible that our more anterior montage engaged only the frontal hub, rather than the entire network, thereby reducing the overall effectiveness of the stimulation.

It is also possible that our stimulation did not reach our target areas - the dmPFC and ACC - due to insufficient dosing. The minimum effective dose of tACS for clear physiological effects has been identified as 0.3 V/m (Wischnewski et al., 2023). Although our stimulation intensity was similar to that of most previous studies, it was less than 0.17 V/m. Compensating for the lower intensity, a longer duration of stimulation and a greater number of sessions could be beneficial, as more sessions are associated with stronger effects (Nissim et al., 2023). However, we note that in Experiment 2, we conducted two stimulation sessions, and the second 25-minute session was not more effective than the first. Future studies might consider increasing the stimulation amplitude to enhance the likelihood of altering the polarization of the underlying neuronal populations and produce noticeable changes in behavioral performance.

Another potential factor is the stimulation frequency (Klink, Pabmann et al., 2020). Different outcomes have been reported depending on the specific conditions: within single study, stimulation at a frequency of 4 Hz over the parietal cortex has been linked to performance improvement, while stimulation at 7 Hz has been associated with performance deterioration (Vosskuhl et al. 2015; Wolinski et al., 2018). Adjusting stimulation frequency to individual theta frequency has yielded negative (Chander et al., 2016), null (Pahor and Jausovec, 2018), or positive (Jausovec & Jausovec, 2014) effects, while decreasing individual theta frequency by 1 Hz positively affected memory performance (Aktürk et al., 2022). As can be seen in our Figure 4a, the natural frequency of theta rhythm was on average 6 Hz, while in the study that used individual frequencies (Jaušovec et al., 2014b; Pahor & Jaušovec, 2018) it lied in the vicinity of 5 Hz. One WM theory suggests that individual memory elements are translated into gamma waves, each embedded in a single theta cycle (Lisman & Idiart, 1995; Lisman & Jensen, 2013). As such, expanding the duration of the theta cycle might enhance WM capacity by allowing more gamma waves within each theta cycle. Although the strongest enhancement of brain rhythms is theoretically expected for tACS waveforms that match the frequency of the targeted endogenous oscillations (Ali et al., 2013; Fröhlich & McCormick, 2010), stimulating endogenous frequencies, as in our study, might not be effective because it does not increase the duration of the theta cycle.

To avoid a potential ceiling effect in memory performance, which could lead to stimulation inefficacy, we used tasks of varying difficulty levels. In Experiment 1, if task difficulty and room for improvement significantly affected the effectiveness of the stimulation, we would expect to observe larger benefits in more complex tasks. For example, accuracy barely exceeded 60% in the alphabetical manipulation of 6 items, compared to around 95% in the simple retention of 4 items. However, despite the large performance differences between conditions, none showed improvements due to the stimulation. In Experiment 2, task difficulty was fixed, but the type of task varied. Most participants did not show perfect accuracy, with an average accuracy of 83.6%. Comparing our results with previous studies, Hu et al. (2022) used a verbal WM task similar to ours, involving the retention of 4 or 6 letters. Their load 4 and load 6 conditions’ accuracy was similar to that in our experiments. They found a significant effect of tACS on RT in the load 6 condition in the high-performance group only. We were unable to replicate this finding. Among other studies showing a moderating effect of task difficulty, Reinhart et al. (2019) used a visual WM task with one item. For young participants the task was easy, and they performed near perfectly, with accuracy well above 90%, while older adults had around 82% performance. Notably, only older adults benefited from the stimulation. In Zeng et al.’s (2022) study, in the n-back task, accuracy was below 80%. The benefits of 8Hz tACS compared with sham were visible in the 3- and 4-back tasks but not in the most difficult 5-back task. In Biel et al.’s (2022) n-back task, results showed above 95% performance across all experimental groups, with no significant effects of tACS on accuracy. We support the notion that the optimal task should strike a balance – difficult enough to avoid ceiling effects, yet not so challenging that accuracy drops to random response levels. However, task difficulty was not a significant moderator in our experiments, suggesting that while this balance may be necessary, other factors also play a large role in determining the effectiveness of tACS.

### Effects of tACS on brain oscillations

Spectral power in the theta frequency band remained unchanged by stimulation during both task and resting state. However, despite setting the stimulation frequency to theta and placing electrodes at the midline, we found a notable decrease in widespread posterior alpha activity during the task’s delay period after verum versus sham stimulation. This effect was absent in resting or not baseline normalized alpha power during the task. This is in contrast to studies of after-effects where theta responded to stimulation (Aktürk et al., 2022; Vosskuhl et al., 2015). Though, we note that two other studies did not find an effect of 6 Hz tACS on theta power (Hsu et al., 2019, 2017). Similar cross-frequency stimulation effects are reported in the literature but also lack consistency. For example, Kleinert et al. (2017) observed non-specific frequency effects of theta stimulation on brain oscillations, showing increased alpha in resting state following verum stimulation, while (Pahor & Jaušovec, 2014) observed non-specific alpha suppression during the task and resting after theta tACS, and Hsu et al. (2017) showed a non-specific increase in beta activity after tACS. Without strong evidence supporting theta tACS’s role in reducing alpha during maintenance of information in WM, it is difficult to draw definitive conclusions.

### Task and load effects on alpha, beta, and theta activity

Our secondary goal was to investigate the general effects related to the type of task and load on oscillatory brain activity. Berger et al. (2014) studied beta, alpha, and theta oscillations during the performance of a similar delayed match-to-sample task involving two types of manipulations and simple retention. We successfully replicated the general effect of the task on theta power, observing a linear correspondence between the level of executive control engagement and theta power (as expressed in the following pattern: alphabetical > backward > forward order conditions). Moreover, tasks that involved retaining and manipulating more memory items also showed an increase in theta power. Altogether these findings support the idea that theta activity is linked to executive control with more complex mental manipulations (Berger et al., 2014; Griesmayr et al., 2010) and higher memory load (Kosachenko et al., 2023) requiring stronger theta engagement.

In the alpha frequency band, Berger et al. (2014) found a pattern of Forward > Backward = Alphabetical, but our results differed, showing Backward = Forward > Alphabetical. Overall, our pattern of results is consistent with the literature that show that alphabetical manipulations lead to a weaker alpha enhancement during the delay (Pavlov & Kotchoubey, 2021). This finding is in line with the inhibition-timing hypothesis (Klimesch et al., 2007) predicting that alpha should gradually release inhibition of sensory cortical areas as access to long-term memory is needed to integrate information from long-term into working memory. The rationale for comparing backward and alphabetical manipulation tasks was that the former does not involve access to long-term memory, whereas the latter does. Thus, the results can be interpreted as alphabetical (semantic) manipulations that require access to long-term memory suppress alpha activity in visual cortical areas. Moreover, we found that an increase in memory load led to an increase in alpha power. This may indicate that tasks not requiring access to long-term memory, enhance alpha activity to suppress irrelevant information that can cause interference to the WM. While this finding aligns with some previous studies, it should be noted that the direction of alpha effects due to increasing load is inconsistent across studies (Pavlov & Kotchoubey, 2022). The conditions in which the direction of the alpha power changes during the delay is important to clarify in the future studies.

Our study found that patterns of lower beta activity were the same as alpha, contrasting with the original study where they differed. Therefore, we could not confirm the proposed relationship between lower beta frequency band and semantic processing that requires long-term memory, as suggested by Berger et al. Importantly, when we followed the original study’s method closely by only considering the 4-letter condition and the first second of the delay period, we observed the disappearance of task effects on both beta and alpha activity.

### Limitations and future directions

The strongest limitation of our study is the degree of control over individual dosage. Variations in individual anatomy may have influenced the amount of current reaching the ACC and dmPFC, potentially rendering the stimulation ineffective in some participants and reducing the overall effect size. Enhancing regional precision and optimizing individual dosage would require a combination of individual electric field modelling and anatomical information obtained from MRI scans. Furthermore, high-density tACS setups could help to make the stimulation more focal, targeting the areas of interest more precisely and avoiding the current spread over larger than necessary brain areas, which can diminish the overall effect. Future research should consider increasing both the intensity above 0.3 V/m to potentially reach effective dosing thresholds and the number of sessions, as well as employing individualized stimulation targets guided by MRI scans. These adjustments could significantly advance the efficacy and precision of theta tACS.

As with other areas of cognitive neuroscience (Garrett-Ruffin et al., 2021; Niso et al., 2022; Pavlov et al., 2021), the tACS literature suffers from a lack of direct replications, preregistered studies, and data sharing (Bikson et al., 2018), and there is direct evidence for publication bias in the field (Héroux et al., 2017). To the best of our knowledge, with one exception (D’Angelo et al., 2023), all preregistered studies and registered reports on healthy populations have failed to confirm predicted effects of tACS on cognition (Biel et al., 2022; Clayton et al., 2018; Fusco et al., 2024; Silas et al., 2023). On the specific topic of tACS for WM addressed in the present study, Biel et al.’s (2022) preregistered study also found no effect of stimulation on accuracy in WM tasks. We encourage future researchers to incorporate practices to improve methodological rigor, including preregistration of hypotheses, analysis plans and make use of registered reports to separate confirmatory from exploratory research and decrease publication bias in this rapidly growing field.

## Data Availability Statement

Behavioral data are publicly available on OSF (osf.io/v2qwc) and EEG data are available on Openneuro (https://openneuro.org/datasets/ds005034).

## Funding Acknowledgements

Author FM is supported in part by a UKRI BBSRC award (BB/X008428/1) and the National Institute for Health and Care Research (NIHR) Leeds Biomedical Research Centre (BRC) (NIHR203331). The views expressed are those of the authors and not necessarily those of the NHS, the NIHR or the Department of Health and Social Care.

## Author Contributions

DK: Investigation; Visualization; Data curation; Formal analysis; Writing - Original draft

OID: Investigation; Supervision

FM: Writing - Review & editing

YGP: Conceptualization; Visualization; Data curation; Formal analysis; Methodology; Project administration; Supervision; Writing - Original draft; Writing - Review & editing

